# Miniature: Unsupervised glimpses into multiplexed tissue imaging datasets as thumbnails for data portals

**DOI:** 10.1101/2024.10.01.615855

**Authors:** Adam J Taylor

## Abstract

Multiplexed tissue imaging can illuminate complex spatial protein expression patterns in healthy and diseased specimens. Large-scale atlas programs such as Human Tumor Atlas Network and are relying heavily on highly-multiplexed approaches including CyCIF and CODEX to image up to 100 antigens. Such high dimensionality allows a deep understanding of cellular diversity and spatial structure, but can provide a challenge for image visualization and exploration. One challenge for data portals and visualization tools is the generation of an informative and pleasing image preview that captures the full heterogeneity of the image, rather than relying on a multi-channel overlay that may be restricted to 4-6 channels. We describe *Miniature*, a tool to automatically generate informative image thumbnails from multiplexed tissue images in an unsupervised and scalable manner. *Miniature* aims to aid researchers in understanding tissue heterogeneity and identifying potential pathological features without extensive manual intervention. *Miniature* uses a choice of unsupervised dimensionality reduction methods including uniform manifold embedding and projection (UMAP), t-distributed stochastic neighbor Embedding (t-SNE), and principal Component analysis (PCA) to reduce on-tissue pixels from a low-resolution, high dimensional image to two or three dimensions. Pixels are then color encoded by their coordinate in low dimensional space using a choice of color maps. We show that perceptually distinct regions in *Miniature* thumbnails reflect known pathological features seen in both the source multiplexed tissue image and H&E imaging of the same sample. We evaluate *Miniature* parameters for dimensionality reduction and pixel color encoding to recommend default configurations that maximize perceptual trustworthiness to both the low-dimensional embedding and high-dimensional image and provide high Mantel correlation between the perceived color difference (delta E 2000) and distance in high- and low-dimensional space. By simulating color vision deficiency, we show that *Miniature* thumbnails are accessible to all. We demonstrate that *Miniature* thumbnails are suitable for a wide range of multiplexed tissue imaging modalities and show their application in the Human Tumor Atlas Network Data Portal.

## Introduction

Highly multiplexed tissue imaging methods continue to demonstrate their power in biomedical applications through their ability to localize multiple proteins of interest (10-100) in high spatial resolution (0.2-1 um/px) and across large regions of interest (500 um - 3 cm) [1]. This family of techniques largely uses antibody labeling of proteins of interest combined with a diverse range of multiplexing methods including fluorophore bleaching (CyCIF [2,3]), antibody stripping (mIHC), DNA probes (CODEX [4,5] and Immuno-SABER [6]), lanthanide mass tags (MIBI [7,8] and IMC [9]) and spectral unmixing (RareCyte Orion [10,11]). They have been extensively applied to reveal new insights into healthy and diseased tissues and are emerging as a future advanced diagnostic tool. As these technologies become more widely adopted and collected at scale, new tools to effectively visualize, select, and triage images through data catalogs and portals are critical to enable researchers to identify datasets of interest, build cohorts, and effectively reuse data.

Frequently, imaging datasets are shared through imaging management platforms such as OMERO [12], repositories such as the BioImage Archive [13], project-specific knowledge portals such as the Human Tumor Atlas Network (HTAN) Data Portal (RRID:SCR_023364) [14] and the HubMAP Data Portal [15,16], or collaborative data platforms such as Synapse (RRID:SCR_006307).

Commonly these platforms present a small image preview or thumbnail to users in a gallery or file view. Image thumbnails have been widely valued as a tool for efficient browsing and selection of images. They provide a condensed and easily viewable representation of an image, allowing for quick scanning and identification of desired images among a large collection. This feature is particularly useful in contexts where a large number of images are present, such as in libraries, galleries, or search results. Implementing image thumbnails can enhance the visual appeal and organization of an image repository or data portal, making navigation and content discovery more user-friendly. Additionally, image thumbnails can serve as a preview mechanism, allowing users to quickly assess the content of an image before opening it in full size.

In particular two tasks, recognition and recall are enhanced and provided by image thumbnails. Users can recognize visual or structural features in a dataset, such as tissue shape or heterogeneity, enabling them to find and select images for further exploration. Users can also recall images that they have previously interacted with, such as finding an image with a tissue of a particular shape or set spatial features. Image thumbnails also add to the visual appeal of a data portal or application, encouraging use and exploration by users. Good image previews for tissue imaging should therefore (1) accurately reflect the spatial heterogeneity and features of the full dataset, (2) be appropriately interpretable for features of user interest and (3) be aesthetically pleasing.

For brightfield or other microscopies resulting in an RGB image, a thumbnail can be simply provided as a downsampled image. Where the file format already contains an image pyramid or specific thumbnail this may be directly extracted from the file. However, for multiplexed tissue more consideration is required. For low-plex images, a multichannel overlay may be used. For three channels, red, green and blue lookup tables (LUTs) may typically be used. For up to six channels cyan, magenta, and yellow may be added to the overlay. Image viewers commonly take this approach. QPath [17] applies unique LUTs ranging in hue, while Napari [18] repeats 6 LUTs (red, green, blue, yellow, cyan, and magenta). While further complementary hues may be added it becomes increasingly difficult for the user to perceptually differentiate different channels. For highly multiplexed images this approach is not therefore effective, resulting in overly bright images with relatively few image features perceivable. This also relies on robust auto-scaling of each channel. Where fewer channels than available in the dataset are used in an image preview, significant information about the sample is not presented to the user. This issue is compounded when the first cycle of an image represents blank, control, or autofluorescence images.

There is therefore an opportunity to provide image thumbnails for multiplexed tissue imaging modalities that capture information present in all channels of the image, reflect tissue features, and are visually appealing. In the broader computer vision field, approaches are available for optimizing thumbnail selection from two-dimensional images, including automatic selection of regions of interest [19]. Strategies also exist for time-series data, such as selecting key frames from a video [20]. However, similar approaches have not been extensively considered or deployed for multi-channel or hyperspectral data.

Encoding higher dimensional data to specific colors has been less frequently considered. Commonly high dimensional datasets including single cell sequencing and multiplexed tissue imaging data (following segmentation and summarisation of marker intensity per cell) are visualized through the use of dimensionality reduction techniques such as PCA, t-SNE and UMAP [21]. Clustering approaches, either in high- or low-dimensional space, or on a kNN graph are frequently used to categorically color cells (or other objects). However, a clustering approach is not optimal for the task of image preview due to the additional steps required in cell-segmentation and quantification.

We looked towards previously described data visualization methods for encoding high dimensional data as colors. Two-dimensional (2D) colormaps have been extensively described [22,23], but underused for encoding coordinates or two variables as colors. 2D colormap selection may be optimized for both elementary localization and identification tasks focussed on single objects or synoptic tasks focussed on comparison of multiple objects [22]. These map strongly to the use of image thumbnails including image re-identification, and assessment of tissue heterogeneity and structure

Previous work in mass spectrometry imaging demonstrated the use of encoding pixels from high dimensional space to colors by representing coordinates in three-dimensional space generated by t-SNE values for red, green and blue in a RGB color space [24]. These approaches proved valuable for exploring MSI data analysis approaches [25] and representing tissue heterogeneity [26]. However the optimisation or limitations of this approach have not been comprehensively explored to date. Previous applications of color encoding high dimensional imaging data include for multispectral UV-vis-NIR imaging using the first three principal components of PCA as red, green and blue [27] ad for dynamic positron emission tomography (dPET), dynamic single photon emission computed tomography (dSPECT) and diffusion tensor magnetic resonance image (DTMRI) encoded as LAB colors [28].

The U-CIE method improves the selection of optimal color encoding for 3D representations of single-cell sequencing data [29]. Instead of simply assigning color axes and orientations, this method optimizes color encoding through rotation, translation, and scaling. This ensures the data fits optimally within the sRGB portion of the LAB color space.

We set out to develop *Miniature*, a tool to generate multiplexed tissue imaging thumbnails using dimensionality reduction and color encoding in 2D and 3D color spaces. We describe its implementation, evaluate its parameter space and optimisation, and demonstrate its applicability to a wide range of multiplexed tissue imaging datasets. We further demonstrate its use within the Human Tumor Atlas Network data portal.

## Methods

### Miniature thumbnail generation

Miniature is designed to work with OME-TIFF images containing an image pyramid [30]. Users may either choose a pyramid level from the image or specify the maximum number of pixels for automatic pyramid level selection. If no pyramid layer with less than the specified maximum number of pixels is available, or the image is not pyramidal, a pyramid is generated, and the appropriate layer is selected.

If a three-channel image is detected, it is rendered as an RGB image, and the output thumbnail is prepared without further action. When enabled, the non-tissue background is removed by summing the channel images, log2 scaling with a pseudo count of 1, and using Otsu’s method to define a threshold. The tissue is assumed to be the brighter portion of the image. This threshold image is cleaned by removing objects smaller than 64 pixels in area. A long array of pixels by channels is prepared from the thresholded multi-channel image. Optionally this matrix is log10 scaled with a pseudo count of 1. If a scaling function is selected this is applied to the array. Scaling options implemented from Scikit-learn include Min Max, Standard Scaling (standardizing features by removing the mean and scaling to unit variance), and Robust Scaling (removing the median and scaling the data according to the interquartile range). Dimensionality reduction is then applied by one of three methods: UMAP, t-SNE, or PCA. For UMAP and t-SNE the distance metric may be selected. The resulting dimensionally reduced array is then used to assign color to pixels in low dimensional space.

Where dimensionality reduction to three dimensions has been used, three approaches for pixel color assignment are provided (RGB, LAB, and UCIE). For RGB pixel color assignment, each dimension is scaled between 0 and 1 with red, green, and blue values assigned to the first, second, and third dimensions respectively. For LAB pixel color assignment the first and second dimensions are scaled between -128 and 127, and the third dimension is between 0 and 100. These values are used to assign the a (green to red), b (blue to yellow), and Lightness components respectively. Where Lab coordinates lie outside the sRGB color space these are clamped to the closest valid value, and converted to a RGB color for display. For UCIE pixel color assignment, we have re-implemented the UCIE algorithm [29] in Python. First, a convex hull is fitted to the low dimensional coordinates. An initial estimate for translation, rotation, and scaling needed to bring the embedded points fully within the hull of the Lab color space is made and applied. Subsequently, the point cloud position with the sRGB portion of L^*^ab color space is optimized by maximizing the size of the point cloud and penalizing points outside the color space through the Nelder-Mead optimization [31]. 1000 iterations are performed for 25 restarts with random rotations of the starting point cloud. Coordinates in the optimized point cloud are then used to assign the L^*^ab color that is converted to an RGB value for display. For dimensionality reduction to two dimensions, pixels are color encoded using a 2-dimensional colormap. The embedding is down-binned to 256 × 256 pixels and the corresponding pixel color is picked from the target colormap image. We implement six 2D colormaps as compared by Steiger et al. [32] and provided by Jäckle (Github:dominikjaeckle/Color2D). To enable repolotting or other reuse, an h5 file containing the background mask, tissue array, embedding, and assigned colors may be output.

### Exemplar data

We evaluated Miniature using open-access multiplexed tissue imaging datasets. *Exemplar-001* and *Exemplar-002* are example datasets from the MCMICRO multiplexed tissue imaging pipeline [33]. *Exemplar-001* is a small lung adenocarcinoma specimen taken from a larger tissue microarray (TMA), imaged using CyCIF with three cycles. Each cycle consists of six four-channel image tiles, for a total of 12 channels. *Exemplar-002* consists of four cores from a TMS imaged by CyCIF with ten cycles for a total of 40 channels. The four cores are two meningioma tumors, one GI stromal tumor, and one normal colon specimen. *CRC04* is an 18-channel RareCyte Orion image of an adenocarcinoma sample. The OME-TIFF image was accessed via Zenodo (Image ID: P37_S32-CRC0) [34]. The five-channel overlay and H&E images were reproduced from a public Minerva story [35]. t-CyCIF, mIHC, CODEX, ImmunoSABER, MIBI and IMC datasets were selected from those available on the HTAN data portal [14].

### Perceptual metrics

We implemented an approach to assess the perceptual trustworthiness of pixel color assignments in relation to their coordinates in both high and low-dimensional space based on existing embedding trustworthiness approaches [36–38]. Here the deltaE2000 color difference formula [39] was used to calculate the distance in perceptual space, euclidean distance was used to calculate the distance in low-dimensional space, and the metric used for the dimensionality reduction was used to calculate the distance in high-dimensional space. Trustworthiness metrics were based on 1000 randomly selected pixels. As trustworthiness provides a measure of how much of the local structure of a data set is preserved we also calculated the Mantel correlation coefficient [40] between perceptual, embedding and original distances to account for a higher scale perceptual preservation.

A combined score was calculated by min-max scaling and summing four metrics: (1) perceptual trustworthiness vs. embedding space, (2) perceptual trustworthiness vs. high-dimensional space, (3) Mantel correlation coefficient between perceptual distance and embedding distance and, (4) Mantel correlation coefficient between perceptual distance and original distance.

### Color vision deficiency simulation

Color vision deficiency simulations of deuteranopia, deuteranomaly, protanopia, protanomaly, tritanopia, tritanomaly, monochromacy, and partial monochromacy were generated using Sim Daltonism 2.0.5 for a *Miniature* thumbnail of the *CRC04* Rarecyte Orion dataset

### Cell segmentation and unsupervised clustering

Cell segmentation of exemplar datasets was performed using MCMICRO [33], using the Unmicst segmentation module [41] with default parameters. Cell-by-feature arrays were further processed with scimap [42]. Leiden clustering was performed with a resolution of 0.2.

### Using Miniature

Usage instructions for Miniature are provided at github.com/adamjtaylor/miniature. The README contains documentation on the use with Python, a Docker container and Nextflow.

## Results and Discussion

### Color encoding of high-dimensional images generate informative image previews

We set out to generate informative and aesthetically pleasing image thumbnails for multiplexed tissue imaging data from a wide range of methods and data contributors. Multiplexed tissue imaging data of any number of channels can be easily presented as a gallery of individual channel images (Figure 1a), but this does not scale well the visualization of very highly multiplexed tissue images or for tissues with many distinct regions defined by unique marker combinations. Many microscopy image file formats contain image pyramids. Image pyramids are a multiresolution representation that efficiently processes images at varying resolutions for tasks like by iteratively downsampling the original high-resolution image into multiple layers with decreasing detail levels. As high-resolution is not typically required of thumbnails we can pull a low-resolution layer of the image pyramid for further processing (Figure 1b). Where an image pyramid is not present, Miniature will appropriately downsample to a smaller image. Using a lower resolution image also facilitates faster dimensionality reduction. Dimensionality reduction approaches may then be used to reduce the multi-channel image to two or three dimensions (Figure 1c) while preserving a level of local or global distance between high and low dimensional space. Miniature supports three common methods: UMAP, t-SNE and PCA, and provides access to a number of parameters for these methods. Colors are then assigned to individual pixels based on their coordinates in low-dimensional space (Figure 1d).

**Figure 1.**
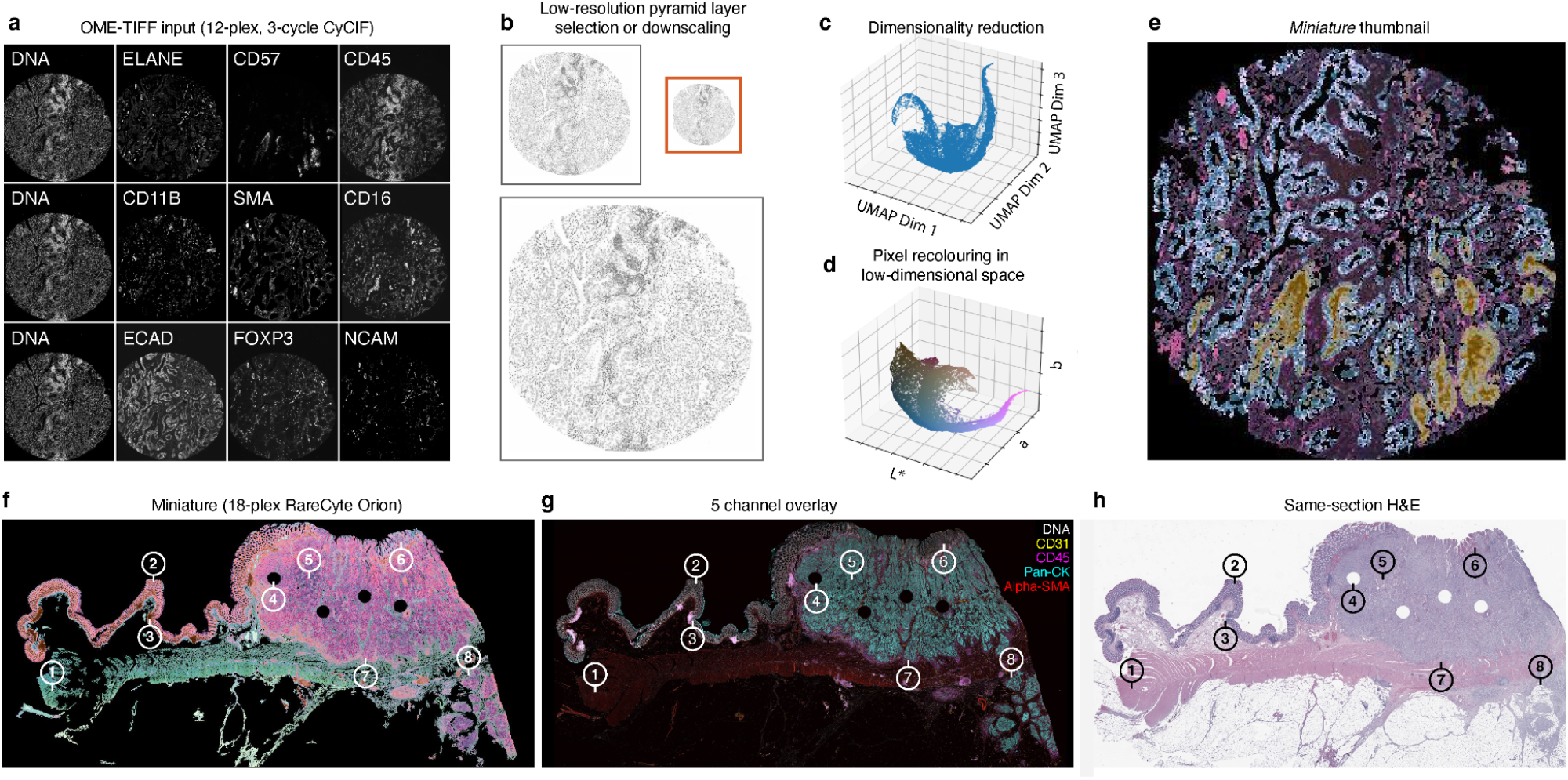
Generation of Miniature thumbnails for informative and aesthetically appealing representation of tissue heterogeneity and pathology. The Miniature technique utilizes unsupervised dimensionality reduction and color encoding to offer comprehensive insights into tissue heterogeneity, serving as a powerful tool for studying gross sample pathology. **(a-e)** Steps involved in creating a Miniature thumbnail from a 12-plex CyCIF image of a TMA core (MCMICRO exemplar-001) using default settings (UMAP to 3D, UCIE colorspace). **(a)** Mosaic of twelve channel images from three cycles representing the OME-TIFF input. **(b)** Selection of a low-resolution layer from the image pyramid. **(c)** Dimensionality reduction to three dimensions using UMAP. **(d)** Recoloring of pixels by optimizing the fit of the embedding point cloud into the sRGB portion of L^*^ab color space with UCIE. **(e)** The resulting Miniature thumbnail image is 314 pixels wide. **(f-h)** Miniature thumbnails exhibit distinct features corresponding to pathological structures. **(f)** Miniature thumbnail generated from an 18-plex RareCyte Orion image of adenocarcinoma (HTA with default settings (UMAP to 3D, UCIE colorspace). **(g)** Five-channel overlay depicting DNA (white), CD31 (yellow), CD45 (magenta), pan-cytokeratin (cyan), and α-Smooth muscle actin (red). **(h)** Hematoxylin and eosin stained brightfield microscopy image of the same section. Highlighted features of interest: **(1)** Submucosa. **(2)** Normal mucosa. **(3)** Peyer’s patch. **(4)** TMA core (1 of 4). **(5)** Invasive adenocarcinoma (submucosal, pT1). **(6)** Entrapped normal mucosa. **(7)** Invasive adenocarcinoma (Invading muscularis, pT2). **(8)** Invasive adenocarcinoma (subserosal, pT3).

### Perceptual recall of tissue heterogeneity and pathology

The resulting Miniature thumbnail (Figure 1e) reveals regions perceptually similar colors that correspond to tissue features seen in the individual channel images (Figure 1a) and Leiden clusters generated from segmented cells (Figure 2f). For the example of a TMA core of colorectal adenocarcinoma imaged by CyCIF, light blue areas clearly correspond to epithelial regions of high ECAD expression and align with Leiden cluster 1, deep purple areas are high in CD45 indicating immune infiltration and align with cluster 0, gold areas are high in CD57 indicating the presence of NK cells and align with cluster 3, and magenta areas are high in SMA, indicating fibrotic regions.

**Figure 2:**
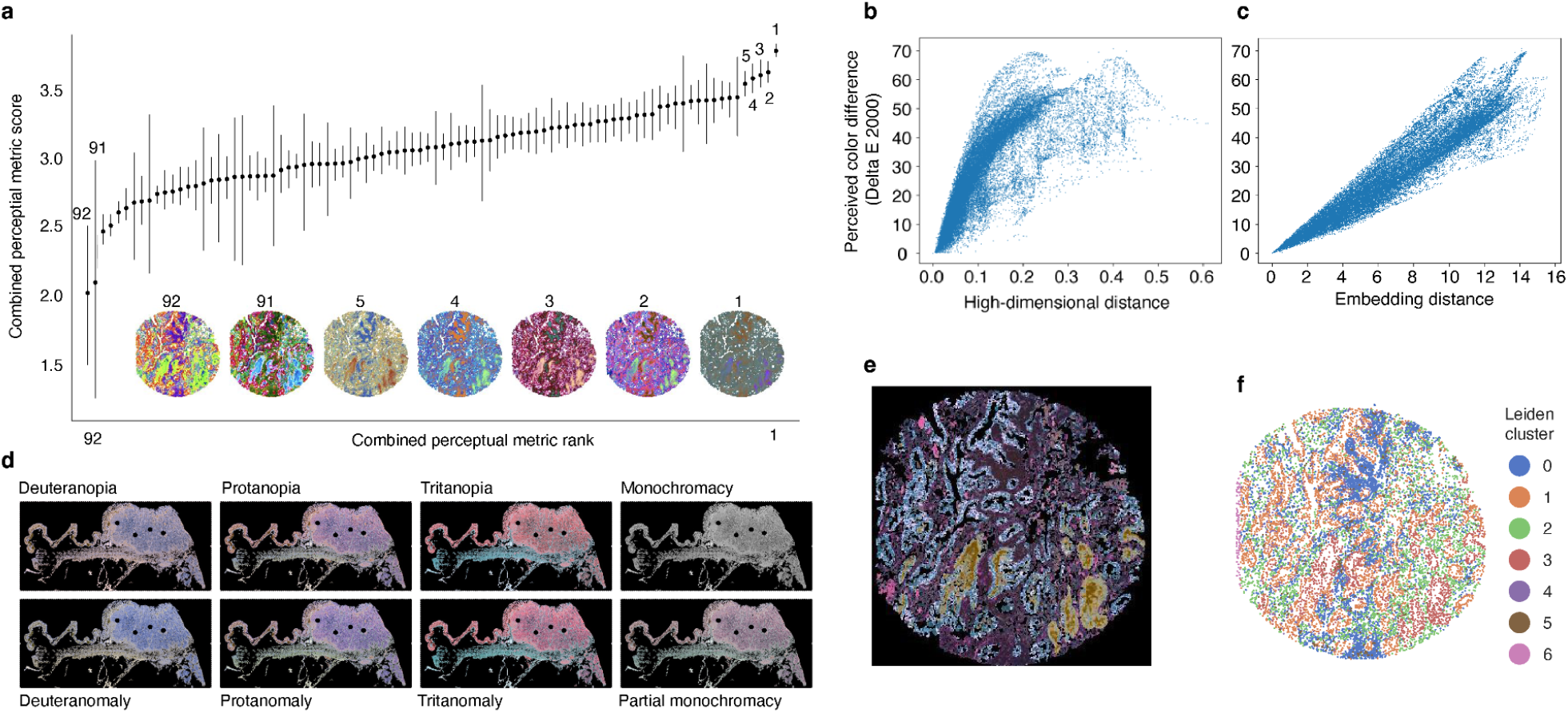
Evaluation of Miniature thumbnails for perceptual trustworthiness, color vision deficiency safety, and representation of segmented-cell clusters. **(a)** Point plot representing the combined perceptual metric score (Sum of the MinMax normalized perceptual trustworthiness and Mantel correlation in high dimensional and embedding space) for combinations of dimensionality reduction method (UMAP, t-SNE, PCA), distance metric (UMAP & t-SNE only: Euclidian, cosine), scaling (unscaled, Scikit-learn RobustScaler) and colormap (32: UCIE, RGB, LAB. 2D: Bremm, cube diagonal, Schumann, Steiger, Teulling 2, Ziegler). Points show the mean score and bars show +/-one standard deviation for Miniature thumbnails generated from five CyCIF images (MCMICRO exemplar-001 and four de-arrayed cores of MCMICRO exemplar-002, max pixels 1E5). Points are ordered by rank. **(1-5, 91, 92)** Miniature thumbnail images from exemplar-001 from selected high- and low-ranked combinations are numbered by their rank. **(1)** PCA to 3D, unscaled, UCIE. **(2)** UMAP to 3D, unscaled, Euclidian, RGB. **(3)** UMAP to 3D, unscaled, Euclidian, UCIE. **(4)** UMAP to 2D, unscaled, Euclidian, cube diagonal. **(5)** UMAP to 2D, unscaled, euclidean, Teuling 2. **(91)** t-SNE to 3D, RobustScaler, LAB. **(92)** t-SNE to 2D, RobustScaler, Ziegler. **(b, c)** Scatter plots depicting **(b)** Euclidian distance in high-dimensional space and **(c)** embedding space versus perceived color difference (delta E 2000) between 1000 randomly selected points from the Miniature thumbnail shown in Figure 2b, subfigure 3 (UMAP to 3D, unscaled, Euclidian, UCIE). **(d)** Color vision deficiency CVD) simulation for the Miniature thumbnail shown in Figure 1f. CVD forms represented are deuteranopia, deuteranomaly, protanopia, protanomaly, tritanopia, tritanomaly, monochromacy, and partial monochromacy. **(e)** Miniature thumbnail of MCMICRO exemplar-001 (UMAP to 3D, unscaled, Euclidian, UCIE, max pixels 1E6). **(f)** Cell segmentation and Leiden clustering (resolution = 0.2) of MCMICRO exemplar-001.

Exploring a Miniature thumbnail generated from a 18-plex RareCyte Orion image of colorectal adenocarcinoma (Figure 1f), we see that features of interest seen in both a selected 5-channel overlay from the multiplexed tissue image (Figure 1g) and same-section H&E image (Figure 1h) that were highlighted in the original publication are visible as perceptually distinct coloured regions in the thumbnail image. These include submucosa (teal in thumbnail), normal mucosa (light orange in thumbnail, Peyer’s patches (maroon in thumbnail) and adenocarcinoma regions (magenta in thumbnail)

### Perceptual trustworthiness and parameter optimization

Encoding high dimensional information as color is potentially fraught with the compounding limitations inherent in dimensionality reduction and human color perception. While the primary goal of *Miniature* was to produce pleasing thumbnails, it was also important to evaluate the perceptual trustworthiness of resulting images in relation to the high-dimensional diversity of the source data. For each combination of dimensionality reduction method, embedding parameter and color encoding method we evaluated both the trustworthiness of the embedding between (high dimensional to low dimensional distance)) and the perceptual trustworthiness of the color encoding (high-dimensional distance to perceptual color difference). Ranking a combined score of perceptual and embedding trustworthiness and Mantel revealed a range of parameter performance (Figure 2a). The highest performing model was PCA to three-dimensions with no scaling and UCIE color encoding (Figure 2a, inset 1). While this had the highest nominal performance the encoded colors were generally not pleasing and often dark. UMAP provided the next 4 highest ranked sets of parameters. With UMAP to three-dimensions with euclidean scaling and RGB (Figure 2a, inset 2) or UCIE color encoding (Figure 2a, inset 3), and UMAP to 2D with *cube-diagonal* (Figure 2a, inset 4) or *Teuling 2* color maps ((Figure 2a, inset 5) all performing strongly and relatively consistently.

Some combinations of parameters had notably lower scores, in particular those using tSNE with RobustScaler The two lowest scoring combinations consisted t-SNE to 3D with RobustScaler and LAB color encoding (Figure 2a, inset 91) and t-SNE to 2D with RobustScaler the *Ziegler* colormap (Figure 2a, inset 92). These do, however, provide bright, colorful and engaging thumbnails that may, where optimal trustworthiness is not required, may be suitable for some applications.

On the basis strong trustworthiness results and visually appeal across a range of modalities and tissue types we initially selected UMAP embedding to three-dimensions with LAB colormap before moving to color encoding with UCIE following its implementation due to its combination of high perceptual trustworthiness while retaining subjectively visually pleasing results.

As seen in a selected example, perceived color difference (Delta E 2000) effectively predicts high-dimensional distance (Figure 2b). As the Delta E2000 value increases, the high-dimensional distance generally rises, demonstrating a positive correlation. For smaller perceived color differences (0.0 to 20.0), the high-dimensional distance shows a rapid increase, while larger perceived differences (beyond 20.0) track with a more gradual increase in high-dimensional distance. This suggests that perceived color difference is a good predictor of high-dimensional distance, although its predictive power slightly diminishes due to increased variability at higher perceived differences.

As expected as color is encoded directly from embedding coordinates, perceived color difference also serves as a strong predictor of embedding distance, exhibiting a nearly linear positive correlation (Figure 2c). As the Delta E 2000 value increases, the embedding distance rises almost proportionally. This indicates that perceived color difference is a highly reliable predictor of embedding distance, with minimal dispersion of data points around the linear trend.

### Accessibility

It is important that image data resources including image previews are accessible and usable by all. Congenital color vision deficiency (CVD) affects as many as 8% of males and 0.5% of females [43]. Using colormaps that facilitate similar interpretation for all regardless of CVD status is therefore an important goal for all scientific visualizations [44]. We simulated major CVD variants for an exemplar Miniature thumbnail of RareCyte Orion data (Figure 2d). While for many CVD variants image contrast is reduced, key attributes of the thumbnail and its effectiveness are retained, including tissue outline and perceived color or contrast differences between major tissue regions.

### Applicability to a multiple modalities in data portals

Multiplexed tissue imaging data are increasingly used alongside single-cell sequencing and spatial transcriptomics as a key assay in multimodal programs generating cell atlases of health and disease. The Human Tumor Atlas Network (HTAN) Data Portal (RRID:SCR_023364, data.humantumoratlas.org) is a web platform designed to host, share, and explore multi-omic cancer data from HTAN atlas programs focussed on generating molecular and cellular atlases of critical transitions in cancer. As of July 2024 the HTAN Data Portal hosts over 2700 multiplexed tissue imaging datasets from eight modalities accross major platforms and vendors. Such data portals are excellent candidates for informative and pleasing thumbnails for multiplexed tissue imaging datasets. Thumbnails help users immediately understand or recognize aspects of the dataset they are about to explore and enable recall and re-identification of known datasets based on image features or expected tissue structures. Miniature thumbnails have been generated for the majority of images released through the data portal and are presented to users both in the file preview page and a larger mouse-over window (Figure 3h). A Nextflow workflow (nf-artist) was developed to scalably and reproducibly automate the generation of Miniature thumbnails and Minerva stories in preparation for display on the HTAN data portal.

**Figure 3.**
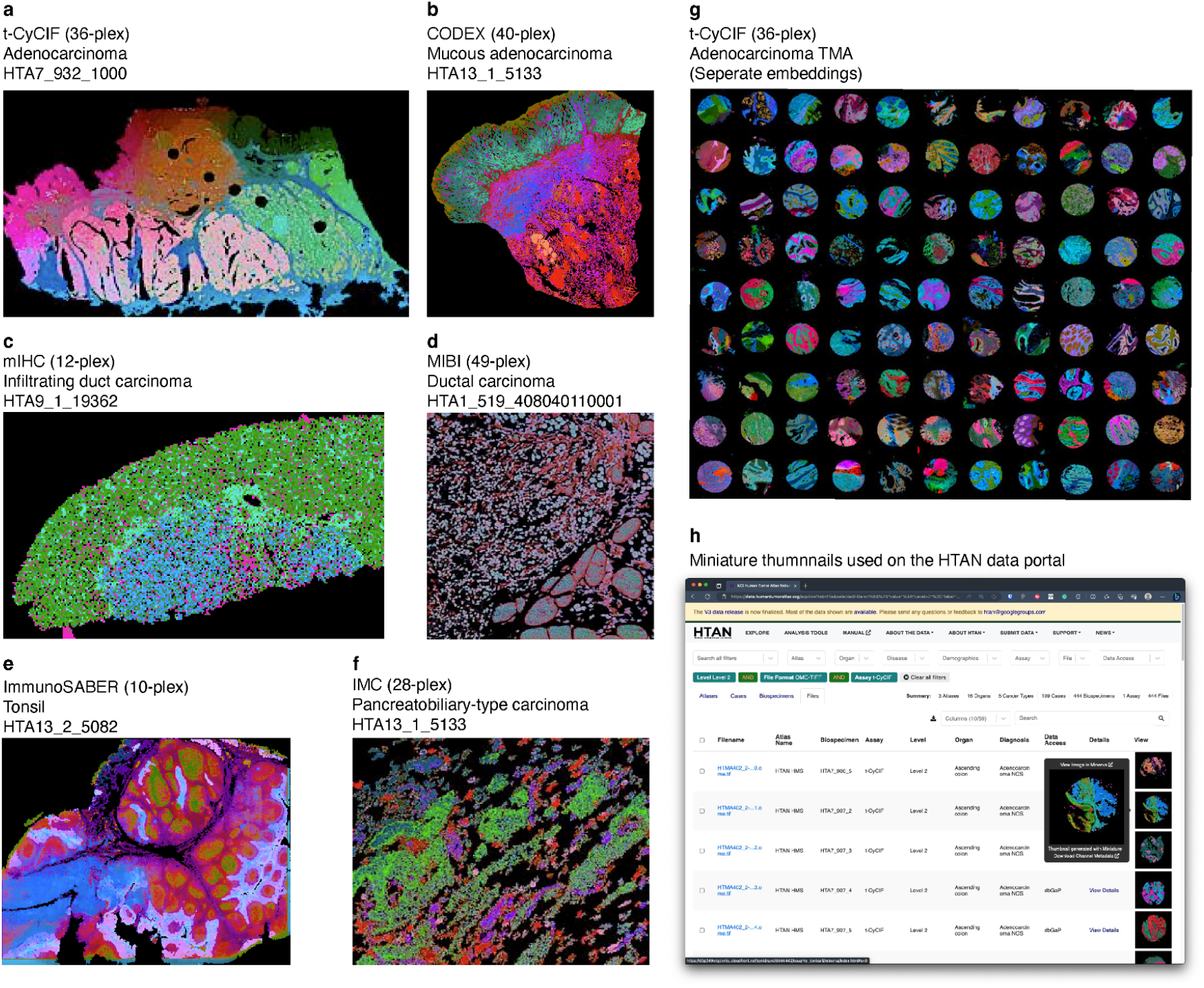
Miniature thumbnails can be generated from a broad range of multiplexed tissue imaging modalities and are ideal for data portals. **(a-g)** Representative Miniature thumbnails of six multiplexed tissue imaging modalities available through the HTAN data portal (UMAP, L^*^ab colorspace, 3D) **(a)** t-CyCIF (36-plex) of Adenocarcinoma. **(b)** CODEX (40-plex) of mucous adenocarcinoma. **(c)** mIHC (12-plex) of infiltrating duct carcinoma. **(d)** MIBI (49-plex) of ductal carcinoma. **(e)** Immuno-SABER (10-plex) of tonsil. **(f)** IMC (28-plex) of pancreaticobiliary-type carcinoma. **(g)** t-CyCIF (39-plex) of a tissue microarray of adenocarcinomas (Miniature thumbnails generated individually for each core. Color spaces are not comparable across cores). **(h)** Screenshot of the HTAN data portal (data.humantumoratlas.org) showing Miniature thumbnails displayed as thumbnails with tooltip expansion.

The use of Miniature thumbnails on the HTAN data portal demonstrate that the dimensionality reduction and color encoding approach taken by Miniature is compatible with a wide range of multiplexed tissue imaging modalities. These include cyclic techniques such as CyCIF (Figures 3a & 3g) and mIHC (Figure 2b), DNA-barcoded antibody methods including CODEX (Figure 3b) and ImmunoSABER (Figure 3e), and microprobe methods using metal-tagged antibodies such as MIBI (Figure 3d) and IMC (Figure 3f). Across these methods Miniature thumbnails demonstrate the attributes of a useful image preview including visualization of tissue shape, along with gradients and discrete features across length scales. These correspond to obvious tissue features such as tonsil germinal centers seen in the preview of a ImmunoSABER image (Figure 3f).

### Limitations and further development

*Miniature* has proved an effective tool for generating multiplexed tissue imaging thumbnails and has been deployed as a part of the Human Tumor Atlas Network Data Portal. There remain several limitations and refinements that may improve its performance and opportunities for use in other imaging tools and databases. While perceptually different colors within miniature thumbnails are demonstrated to closely relate to known clusters it is not possible to show which channels contribute to these regions, unlike a more typical multi-channel composite. A potential approach may be to develop semi-automated approaches to identify three to six channels that are most representative of clusters and generate composite images for these. This would however not represent the full heterogeneity of very high dimensional datasets. In many cases multiplexed tissue imaging datasets contain multiple channels (often entire cycles) made up of blank, antibody-control or autofluorescence channels. Depending on the scaling parameters used Miniature may overemphasize low-intensity features such as uneven illumination or tiling within these channels that may interfere with information arising from tissue heterogeneity in channels with positive staining. Dimensionality reduction approaches including UMAP and t-SNE can be computationally expensive in comparison to simple multi-channel overlay, even for relatively small thumbnails which may contain more than 50k pixels. *Miniature* may therefore not be suitable for applications where on-the-fly generation of thumbnails is required and is better suited for pre-rendering of resources.

Miniature is complementary to and uses elements of tools to encode color in two and three dimensional scatter plots including U-CIE, and is complementary to approaches such as Mystic [45], which enable comparison between collections of images in low-dimensional embeddings.

*Miniature* may benefit from extension to encompass further dimensionality reduction methods such as SOMs or autoencoders, and enabling the use of additional or custom colormaps. Similar approaches for color encoding of embeddings are likely applicable and useful for various multiplexed spatial assays beyond antibody-based imaging, particularly in situ sequencing and spot-based spatial transcriptomics, such as 10x Xenium, Nanostring CosMX, and Slide-seq. In the future Miniature-like thumbnails may be useful reporting outputs from multiplexed tissue imaging workflow such as MCMICRO or as available rendering states or overlays in visualization tools including Minerva, Vittessve or Napari.

## Conclusions

The development and implementation of *Miniature* offer a robust and effective approach for generating informative and aesthetically appealing thumbnails for multiplexed tissue imaging datasets. This tool employs unsupervised dimensionality reduction techniques coupled with a range of color encoding methods, to visualize complex tissue heterogeneity and pathological features. Our findings demonstrate that Miniature effectively captures and represents the intricate details of high-dimensional tissue images, making it easier for researchers to explore and select relevant datasets. The perceptual trustworthiness of the thumbnails was evaluated, revealing that UMAP to three dimensions with UCIE color encoding provided the best combination of visual appeal and accuracy in representing the high-dimensional data. The tool was shown to produce perceptually distinct regions in the thumbnails that correspond to known pathological features, as evidenced by the comparison with segmented cell clusters and traditional H&E staining. The application of Miniature across various multiplexed imaging modalities, including CyCIF, CODEX, mIHC, MIBI, Immuno-SABER, and IMC, highlights its versatility. Its integration into the Human Tumor Atlas Network Data Portal further demonstrates its practicality, providing users with immediate visual insights into the structure and heterogeneity of tissue samples. Additionally, simulations for color vision deficiencies ensured that the generated thumbnails are accessible to a broader audience. Overall, Miniature represents an advancement in the visualization of multiplexed tissue imaging data, facilitating rapid recall and assessment of tissue architecture and pathology.

## Data and code availability

Code, documentation, and examples of use are available on GitHub. (https://github.com/adamjtaylor/miniature/). Instructions for accessing CyCIF images Exemplar-001 and Exemplar-002 are available on the MCMICRO website (mcmicro.org). HTAN datasets are available via the HTAN Data Portal (data.humantumoratlas.org).

## Acknowledgements

The authors gratefully acknowledge Jeremy Muhlich, Amanda Kwong, Dev Soni, Jie Zohu, Mark Keller, and Surya Narayanan Hari for their contributions to developing the idea of multiplexed image preview representation during the CSBC/PS-ON Image Analysis Working Group Hackathon. We thank Clarence Yapp, Peter Sorger, Ashley Clayton, Alex Lash and members of the HTAN Data Coordinating Center for their helpful discussions and comments.. We appreciate Ino de Bruijn for integrating Miniature thumbnails into the HTAN data portal, and Thomas Yu, Rixing Xi, and Brad Macdonald for helping with the integration of Miniature into a Nextflow pipeline. We also thank Alex Dexter, Alan Race, and Josephine Bunch at the National Physical Laboratory for discussing and developing foundational concepts in high-dimensional spatial data visualization, which inspired and informed this work. This research was funded by the National Cancer Institute as part of the HTAN Data Coordinating Center (U24CA233243).

